# Detection of Activated Receptor Tyrosine Kinases in Human Lung Squamous Cell Carcinoma

**DOI:** 10.1101/2023.02.07.527536

**Authors:** Nancy Kendrick, Matt Hoelter, Andrew Koll, Costel C. Darie, Jon Johansen

## Abstract

Lung squamous cell carcinoma (LSCC) has a high mutational burden and poor prognosis, even with immunotherapy. In the Lux-Lung 8 trial, afatinib, an epidermal growth factor receptor (EGFR) inhibitor, showed a long-term benefit in 5.3% of patients with LSCC. Because activating mutations of EGFR are rare in LSCC, the response was likely due to wild-type EGFR being activated by an unknown mechanism. All receptor tyrosine kinase (RTK) proteins, both wild-type and mutated, are activated by phosphorylation of specific tyrosines which serve as binding sites for various SH2 proteins. The aim of this feasibility study was to determine whether enhanced chemiluminescent western blotting (WB) with a phosphotyrosine (pTyr) antibody is sufficiently sensitive to detect pTyr-RTK proteins in human LSCC tissues. We performed WB analysis on 25 resected human lung tissue samples, including 12 LSCC, two adenocarcinomas (LADC), and 11 control (non-tumor) lung samples. The analysis showed ∼220 kDa pTyr-protein bands in two LSCC samples that were much more abundant than the corresponding bands in controls or LADC samples. To identify pTyr-RTKs, pTyr WB patterns of the two samples were compared to those of five RTK candidates: EGFR, platelet-derived growth factor receptor beta (PDGFRB), vascular endothelial growth factor receptor, anaplastic lymphoma receptor, and mesenchymal-epithelial transition factor. The strong pTyr signal in one sample matched EGFR, whereas the other matched a combination of EGFR and PDGFRB. We conclude that pTyr WB is sufficiently sensitive to detect pTyr-RTK drivers in flash-frozen tumor tissues and might identify LSCC patient subsets responsive to RTK inhibitors.

## Introduction

Lung cancer is the leading cause of cancer-related deaths in the United States, with over 135,000 deaths annually. Approximately 85% of lung cancers are non-small cell lung cancer (NSCLC), comprising mainly lung adenocarcinoma (LADC, ∼40%) and lung squamous cell carcinoma (LSCC, ∼25%) [1, 2]. Although the mean mutation rates of LSCC and LADC are similar, 8.7 and 9.7 mutations/Mb respectively [3-5], the two cancers are surprisingly different. The mean doubling time of LSCC is much shorter, 126 ± 58 days, compared to 976 ± 3134 days for LADC [6]. The 5-year overall survival (OS) rate for LSCC is shorter at every stage [7]. Differences may stem from the cells of origin: LADC arises primarily from cells that secrete surfactant components, whereas LSCC originates from cells that line the inside of the lung airways [2].

Precision oncology [8, 9] is a personalized approach for cancer treatment in which therapeutic drugs are matched to genomic biomarkers detected in a patient’s tumor. Patients with LADC are more likely than those with LSCC to show actionable receptor tyrosine kinase (RTK) biomarkers detected by Next Generation Sequencing (NGS). Activating mutations of the epidermal growth factor receptor (EGFR) are present in 15-20% of LADC patients and of the anaplastic lymphoma receptor (ALK) in 5-7%. Inhibitor drugs targeting mutated EGFR and ALK proteins show initial LADC patient response rates of approximately 70% and 60%, respectively [10, 11]. In contrast, few actionable mutations of EGFR or ALK are present in LSCC tissue [3, 5].

In the LUX-Lung 8 trial, in which EGFR inhibitors afatinib and erlotinib were compared as second-line treatments for late-stage LSCC patients, a small subset of patients in both groups showed long-term benefits. For afatinib, 21/398 patients (5.3%) achieved a median overall survival of 34.6 months[12]. Both WT and mutant EGFR are inhibited by afatinib [13]. However, the value of administering afatinib to all LSCC patients eligible for second-line treatment when only ∼5% will respond is debatable [14].

NGS genomic profiling to identify mutated drivers has become routine, but at least half of the patients who undergo tumor genomic profiling have no actionable alterations. Transcriptome profiling is complex and is not yet in place [15]. A method to identify actionable RTK drivers is urgently needed.

RTK activity is highly regulated in tissues [16]. EGFR protein, for example, remains inert until one of its seven ligands [17] epidermal growth factor for example, binds to an external domain. Ligand binding triggers EGFR protein dimerization and transphosphorylation of multiple tyrosine residues on the cytosolic side of the membrane, which in turn serve as binding sites for critical cytosolic proteins. Thus, phosphotyrosine (pTyr) residues and surrounding amino acids become docking sites for Src Homology domains (SH2) on cytosolic proteins [18] that relocate and interact to trigger cell growth [16]. RTK proteins are activated by tyrosine phosphorylation in the sense that they remain inert without it and bring about cascades of downstream processes once it has taken place. RTK gene amplification, mutation, or protein overexpression are not predictive of inhibitor drug response [19]. Tyrosine phosphorylated RTK proteins are the actual drug targets [20].

However, it is difficult to screen for the latter. Mass spectrometry (MS) is commonly used to identify thousands of proteins simultaneously but cannot distinguish between intact pTyr-RTKs and degraded fragments. Immunohistochemistry (IHC) with a general pTyr Ab cannot distinguish between pTyr-RTKs and pTyr-protein substrates of Src and other TKs; there are too many Ab choices for specific pTyr-RTK residues on multiple candidates. We suggest that an alternative protein analysis method, pTyr WB, could be used to screen human research biopsies to identify those with active pTyr-RTKs.

Here, we describe the results of a feasibility study to determine whether ECL WB is sensitive enough to identify pTyr-RTKs in LSCC biopsy samples using a validated pTyr Ab. A strong pTyr-RTK signal was observed in some LSCC samples, but not in the control samples, as would be expected from a cancer driver. This suggests that WB might be useful for the unequivocal identification of cancer biopsies with specific pTyr-RTK drivers. Characterization of the latter using multiple methods including MS and mRNA fingerprinting might guide precision medicine to better predictions for RTK inhibitor drugs.

## Results

### Samples

We purchased 14 resected human lung tumors and 11 control lung samples from a human biobank in two groups, as shown in Table 1. The NSCLC tissues included 12 LSCC and two LADC tumors. Normal adjacent tissue (NAT) controls were obtained for eight LSCC and one LADC tumor, along with two unmatched non-cancerous lung disease controls, asthma, and tuberculosis.

**Table 1.**
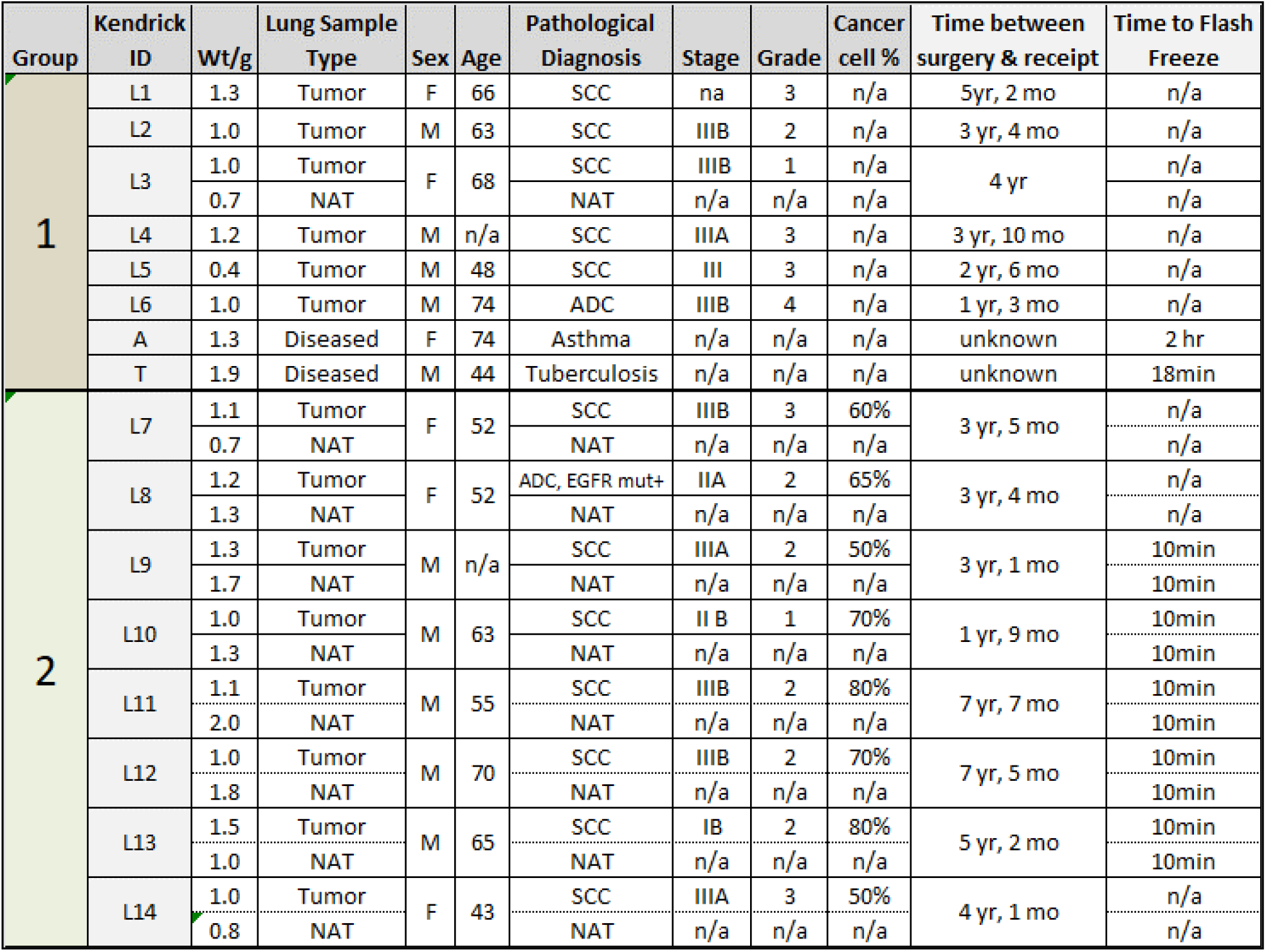
Sample identification for 14 lung tumor samples, 9 matched normal adjacent tissue samples (NAT) and 2 non-cancer diseased lung tissue samples, asthma and tuberculosis, used as negative controls. Group 1 samples were purchased in 2011. Group 2 samples were purchased from the same biobank in 2014. n/a = not available.

## 1D pTyr WB (1D WB)

Figure 1 shows a 1D pTyr WB loaded with the Group 1 samples described in Table 1, including five LSCC, one LADC, and three controls. Equal protein was loaded (40 μg/lane), as recommended by the Journal of Biological Chemistry Editorial Board for the normalization of WBs [21]. Comparison of the LSCC bands to those of the controls revealed that three tumor samples, L3, L4, and L5, had pTyr bands at ∼220 kDa that were darker than those of any of the controls or the other three tumors. The L3 and L5 pTyr bands were pronounced; the L4 sample band was fainter but still more prominent than that of the controls. Additional fainter pTyr bands between 14 and 94 kDa were present and varied between the samples.

**Figure 1.**
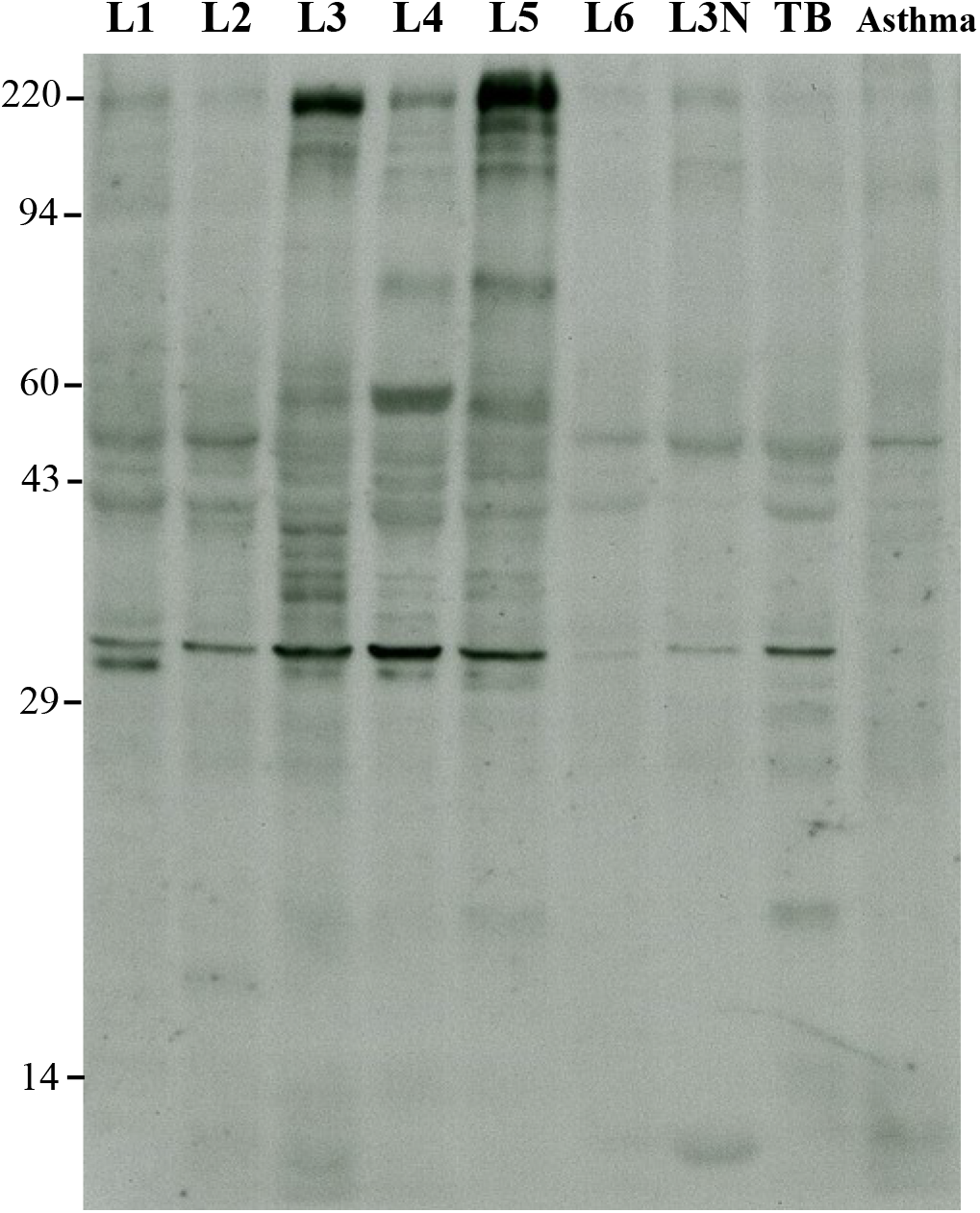
Phosphotyrosine WB pattern for five LSCC (L1-L5), one LADC (L6), and three lung control samples including L3 normal adjacent tissue (L3-N), tuberculosis and asthma samples. All samples were identically prepared by thawing and homogenization on ice in SDS buffer minus beta-mercaptoethanol (BME) with protease and phosphatase inhibitors. After homogenization, the tubes were heated in a boiling water bath until clarification, protein concentrations measured, and BME added. Each prepared sample was divided into multiple single-use aliquots and stored at -80° C. The dark bands in L3, L4, and L5 running at ∼220 kDa are presumptive pTyr-RTK bands.

RTKs are large transmembrane proteins containing multiple tyrosine residues that, once phosphorylated, contribute to oncogenic cell growth [22]. The pTyr-protein bands at ∼220 kDa were likely RTK proteins. But which ones?

Fibroblast growth factor receptor 1 (FGFR1) is amplified in LSCC but has a theoretical molecular weight (MW) of 92 kDa, which is too low to match the ∼220 kDa band [23]. Another possibility, the discoidin domain receptor tyrosine kinase 2 (DDR2), also has a low MW of approximately 120 kDa [24]. Higher MW RTKs implicated as LSCC drivers include EGFR, vascular endothelial growth factor receptor (VEGFR), anaplastic lymphoma receptor (ALK), mesenchymal-epithelial transition factor (MET) and platelet derived growth factor receptor beta (PDGFRB) [25, 26].

Attempts to directly identify the protein(s) co-migrating with the pTyr-RTK signal in tumor L5 by MS failed, as shown in Supplemental Figure S1. Rikova et al. showed that MS identification of pTyr-RTKs from trypsin digests requires the concentration of phosphorylated peptides by affinity resins [27], which is beyond the scope of this exploratory project.

### Identification by 1D matching eliminated MET, VEGFR, and ALK

To determine whether the pTyr signal co-migrated with any of the candidates, we probed identically loaded 1D gels with commercial antibodies against the five candidate RTK proteins. Figure 2 shows the positive results. The pTyr WB from Figure 1, cropped to show the 220 kDa region, is shown for comparison. The VEGFR-2 and ALK WBs were very faint or blank in this MW region after long exposures (not shown).

**Figure 2.**
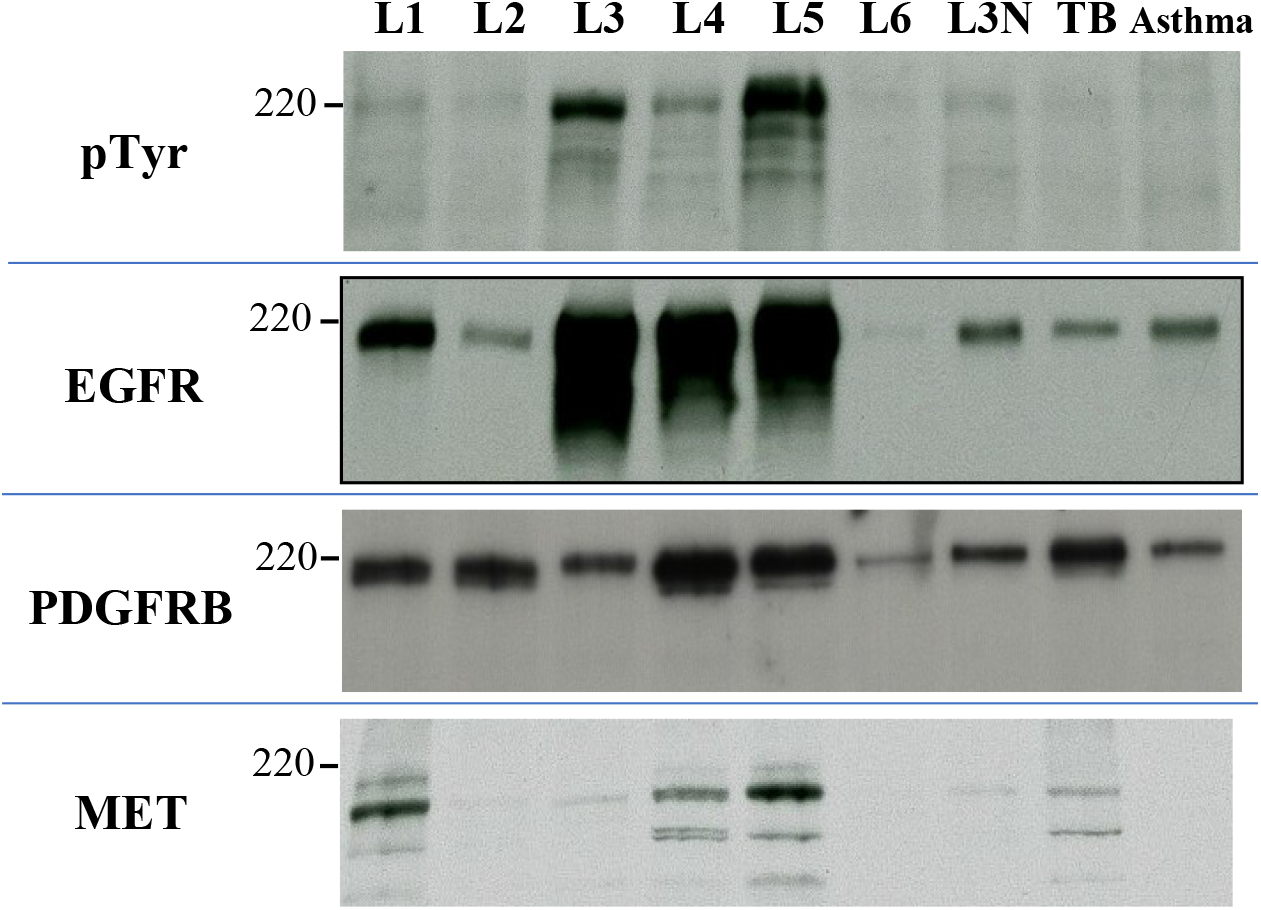
WB images from four 1D gels identically loaded with 40 μg protein each from five SCC tumors (L1-L5), one ADC tumor (L6) and three non-cancerous lung controls: L3-N where N = normal adjacent tissue, tuberculosis (TB), and asthma. The WBs were probed with the indicated antibodies.

The pTyr bands comigrate with both the EGFR and PDGFRB WB bands in LSCC tumors L3, L4, and L5, suggesting that EGFR, PDGFRB, or both were activated to different degrees by tyrosine phosphorylation. MET WB revealed multiple protein bands with the largest migrating at only ∼150 kDa, which is too small to match the pTyr band.

As shown in Figure 3, Group II tumor samples from Table 1 and matching normal adjacent tissue (NAT) controls showed less dramatic pTyr-protein bands at ∼220 kDa compared to bands in freshly thawed aliquots of L3, L4, and L5 run as positive controls. Sample L11 showed a 220 kDa doublet at the intensity of L4, which was slightly greater than that of any of the control samples. In general, the pTyr band intensity was not correlated with the EGFR band intensity.

**Figure 3.**
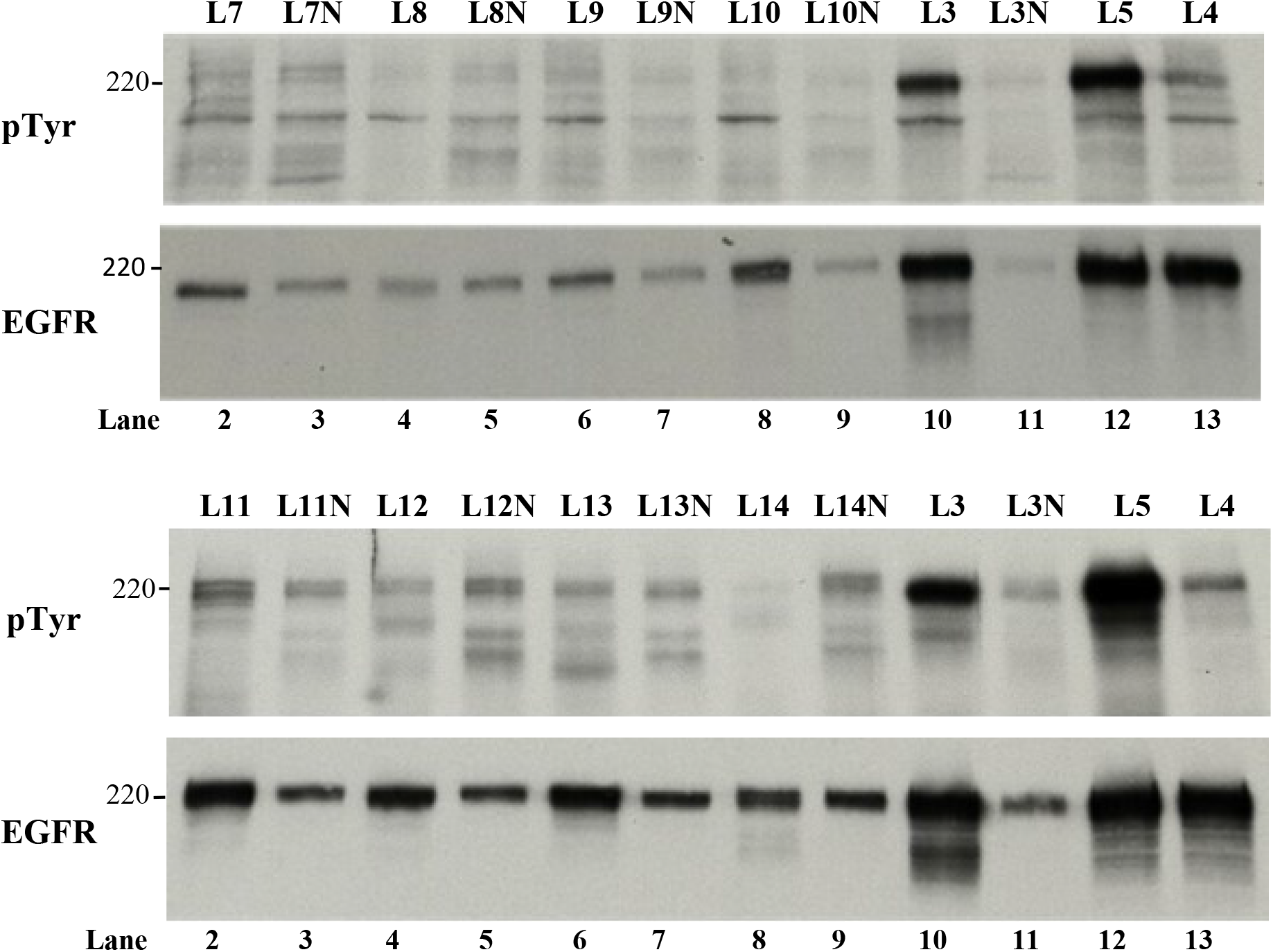
Western blot patterns from seven LSCC and one ADC tumor (L8, L8N), each with matching NAT controls (N). Samples L3, L3N, L5 and L4 were run in the last four lanes on every gel as controls to verify method reproducibility.

Sample L8, a LADC tumor with an EGFR exon 19 activating mutation, was purchased as a positive control with the expectation that it would show a strong pTyr WB band that matched an EGFR band. However, L8 showed faint WB bands for both EGFR and pTyr compared with the strong bands for the L3 sample run on the same 1D gel.

Since Group II samples showed relatively light pTyr bands, we focused on the unequivocal identification of the strong bands in samples L3 and L5. Are these two phospho-proteins EGFR as might be expected from overexpression, or something else?

## Identification of the 220 kDa pTyr-proteins using 2D SDS PAGE

The EGFR and PDGFRB bands in L3 and L5 were detected at the same MW as the pTyr band, so 1D WB does not suffice. Since there were multiple RTK candidates and multiple pTyr antibodies for each of them, identification using specific pTyr antibodies was too costly. Instead, we used an alternative method, 2D SDS PAGE WB (2D WB), with the expectation that adding the orthogonal step of isoelectric focusing (IEF) would facilitate matching. Since the IEF system used in our lab for 2D is compatible with SDS, 1D/2D patterns correlate well [28].

Figure 4 shows pTyr 2D WBs obtained from the six tumor and three control samples shown in Figure 1. Each gel was loaded with 200 μg of tumor lysate, five times the 1D load. The samples give complex 2D protein patterns when a sensitive silver stain is used (Supplemental Figure S2), but only a few pTyr-proteins are detected by pTyr WB.

**Figure 4.**
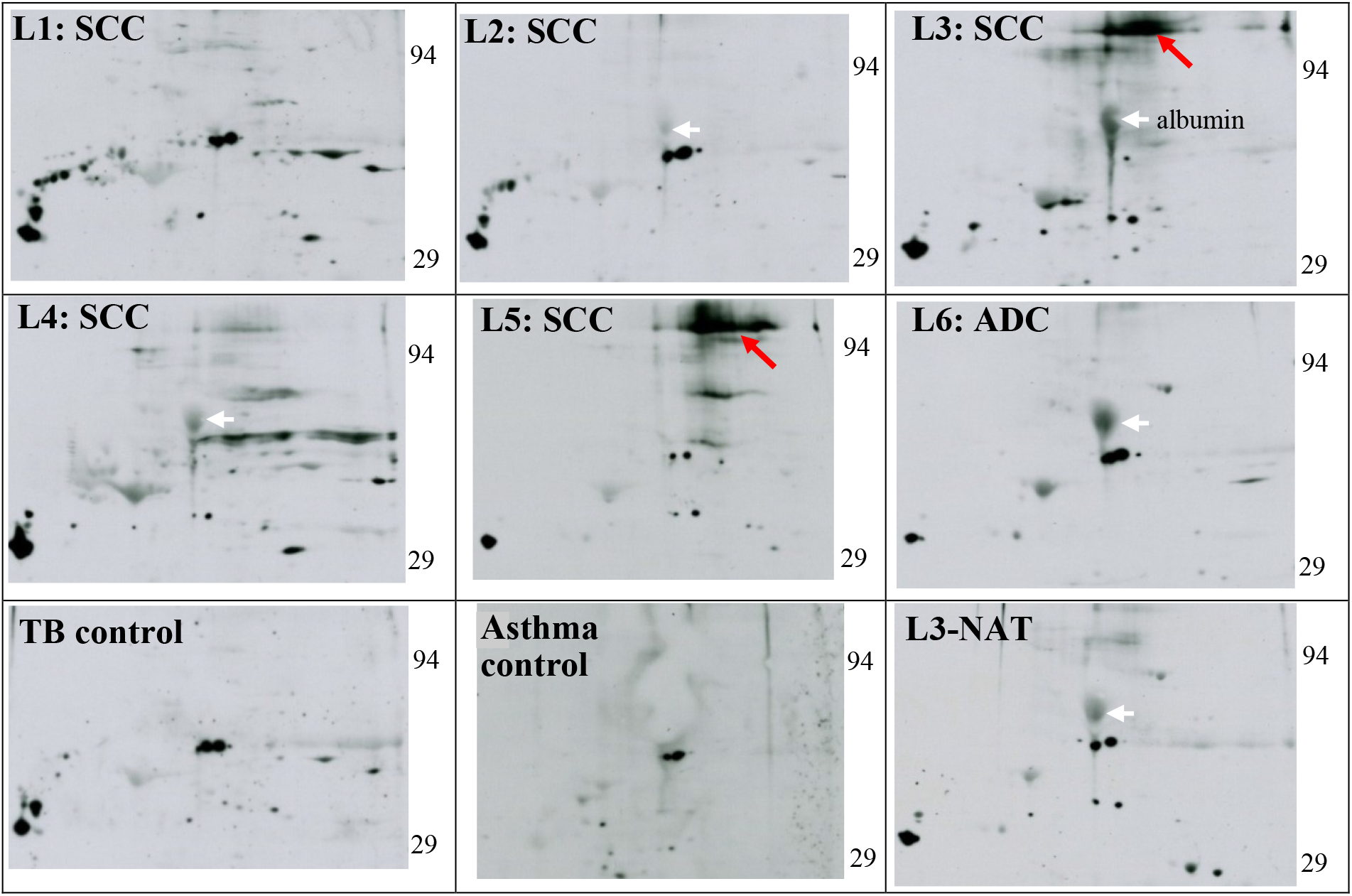
Nine 2D-SDS PAGE pTyr WBs loaded with 200 μg each of Group 1 samples. Two of the six LSCC samples, L3 and L5, showed strong pTyr-protein spots at ∼220 kDa with irregular borders suggestive of heavy glycosylation (red arrows). L4 showed a similar fainter signal at that MW. Serum albumin (white arrows), an abundant protein visible on Coomassie stained PVDF membranes, sometimes stains nonspecifically on WBs and serves as an internal marker.

The strong diffuse ∼220 kDa pTyr-protein spots present in L3 and L5 (red arrows) correspond to the pTyr bands observed in the 1D WB (Figure 1). The diffuse pattern of the pTyr spots is indicative of charge heterogeneity due to heavy glycosylation.

### Identification of the high MW pTyr-protein RTKs using 2D pattern matching

A sensitive silver stain showed only a faint haze in the high MW pTyr area of a 2D gel loaded with 50 μg of L5 (Supplemental Figure S2), indicating the pTyr-protein was in low abundance. Since the tumor samples were prepared in SDS buffer with protease and phosphatase inhibitors to maximize intact protein recovery, the samples were unsuitable for the affinity resin concentration of phosphopeptides required for MS [27]. Thus, an alternative approach was used to compare the 2D pTyr WB pattern with those obtained using anti-EGFR and anti-PDGFRB antibodies on the same samples to look for co-migration.

Figures 5 shows results for samples L3, L4 and L5. The major area of the EGFR WB matches the pTyr WB for all three samples. The L3 PDGFRB WB matches the L3 pTyr WB poorly. The L4 dark pattern is skewed left but a portion of the right side overlaps the faint pTyr signal. However, the entirety of the L5 PDGFRB WB signal falls within the L5 pTyr WB signal. Some pTyr-protein spots (red arrows) do not match to either EGFR or PDGFRB.

**Figure 5.**
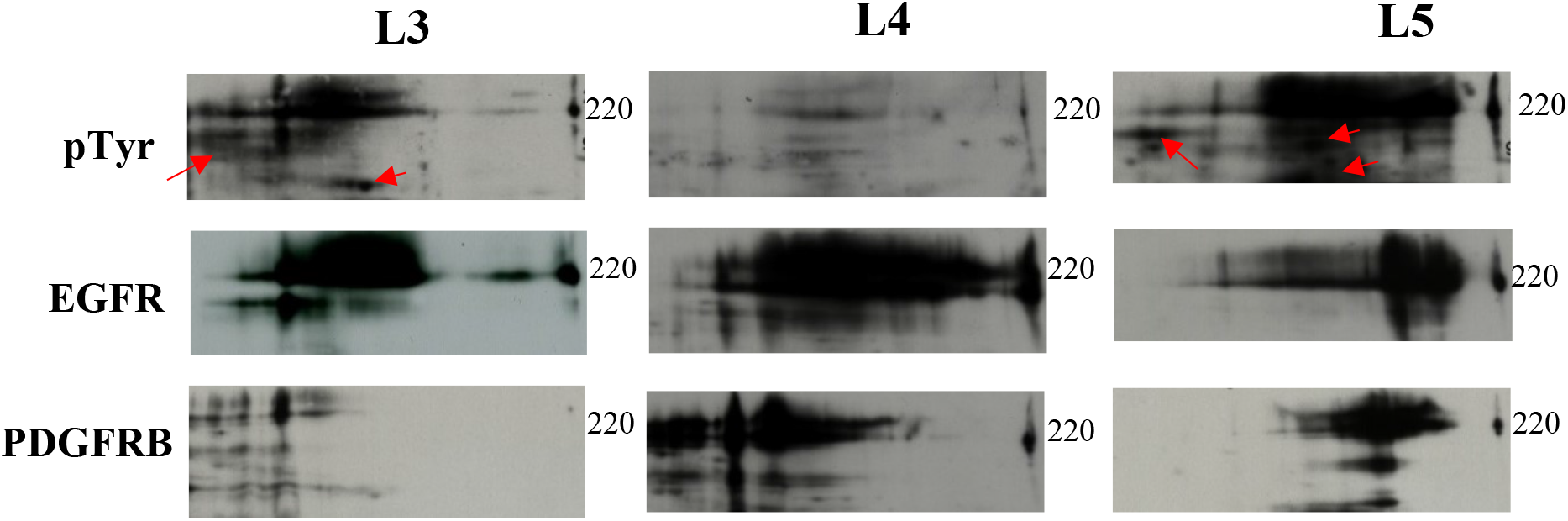
pTyr, EGFR and PDGFRB 2D WB comparisons of tumor samples L3, L4 and L5. Each sample (200 ug) was identically loaded on three 2D gels for WB with antibodies against pTyr, EGFR, and PDGFRB, i.e., 1 antibody per gel, 9 gels total. Red arrows indicate pTyr protein spots that do not match to either.

Heavy RTK glycosylation, in combination with 2D gel-to-gel variation, causes uncertainty in pattern matching. To achieve unequivocal matching, considerable effort was put into strip-and-reprobe experiments, where WBs are performed on the same membrane and thus aligned perfectly. However, in practice, the strip-and-reprobe method is problematic. Re-probed WB images sometimes show spots for unknown reasons. Thus, each re-probed image had to be checked by comparison with an unstripped image from another gel and disallowed if artifacts were present. Despite technical limitations, the strip-and-reprobe results were convincing in some cases (Figure 6).

**Figure 6.**
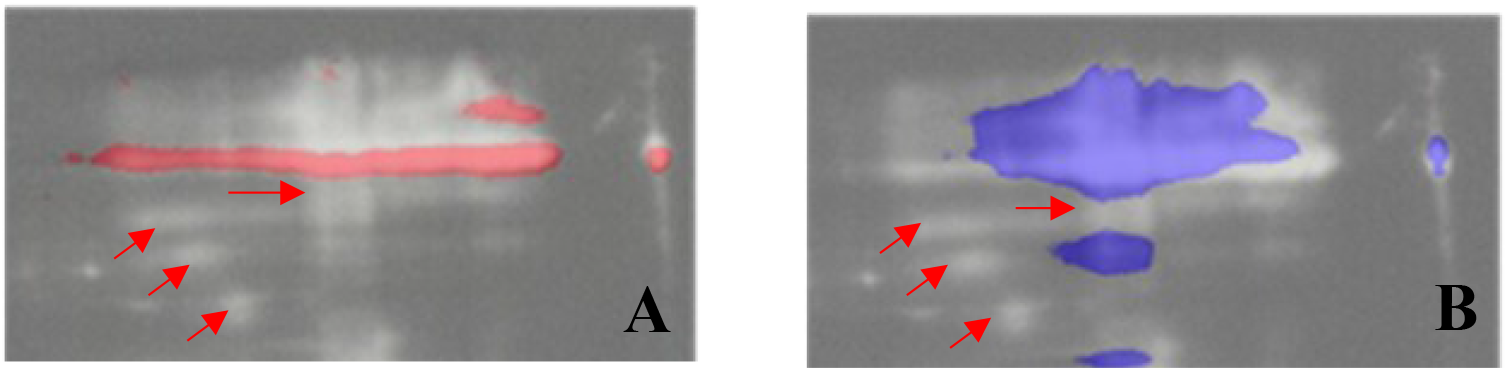
Double strip-and-reprobe experiment for sample L5. A single 2D PDGFRB membrane from sample L5 was blotted with three different antibodies in the order PDGFRB/strip/pTyr/strip/ EGFR. The pTyr WB image was inverted to give a white pattern on a black background using Adobe Elements software. Spots on the EGFR WB image were colored red while those on the PDGFRB image were colored blue. Finally, the RTK images were superimposed over the pTyr image. A: Red EGFR pattern over the inverted white pTyr image; B: Blue PDGFRB over the white pTyr pattern. The cropped PDGFRB WB for L5 is also shown in Figure 5. Red arrows show pTyr spots that don’t match either EGFR or PDGFRB.

This strip-and-reprobe figure shows that a mixture of pTyr-EGFR and pTyr-PDGFRB accounts for most of the pTyr WB signal in L5. Convincingly, at the IEF origin area on the right, the red EGFR colors the bottom half of the origin spot, while the blue PDGFRB superimposes with the upper half, suggesting co-migration with different pTyr-protein isoforms. Some of the lower MW white pTyr signals (red arrows) remain unmatched, as was also observed in Figure 5.

### PDGFRB in LSCC tumor L5 breaks down differently than in L1-L4

Multiple PDGFRB protein spots were detected by 2D WB of LSCC tumors L1-L5 (Figure 7). For L1, L2, L3, L3-NAT and L4, the PDGFRB WBs show complex patterns suggestive of breakdown products from endocytosis with lysosomal degradation [29]. However, for L5, the pattern was simpler and different from the others.

**Figure 7.**
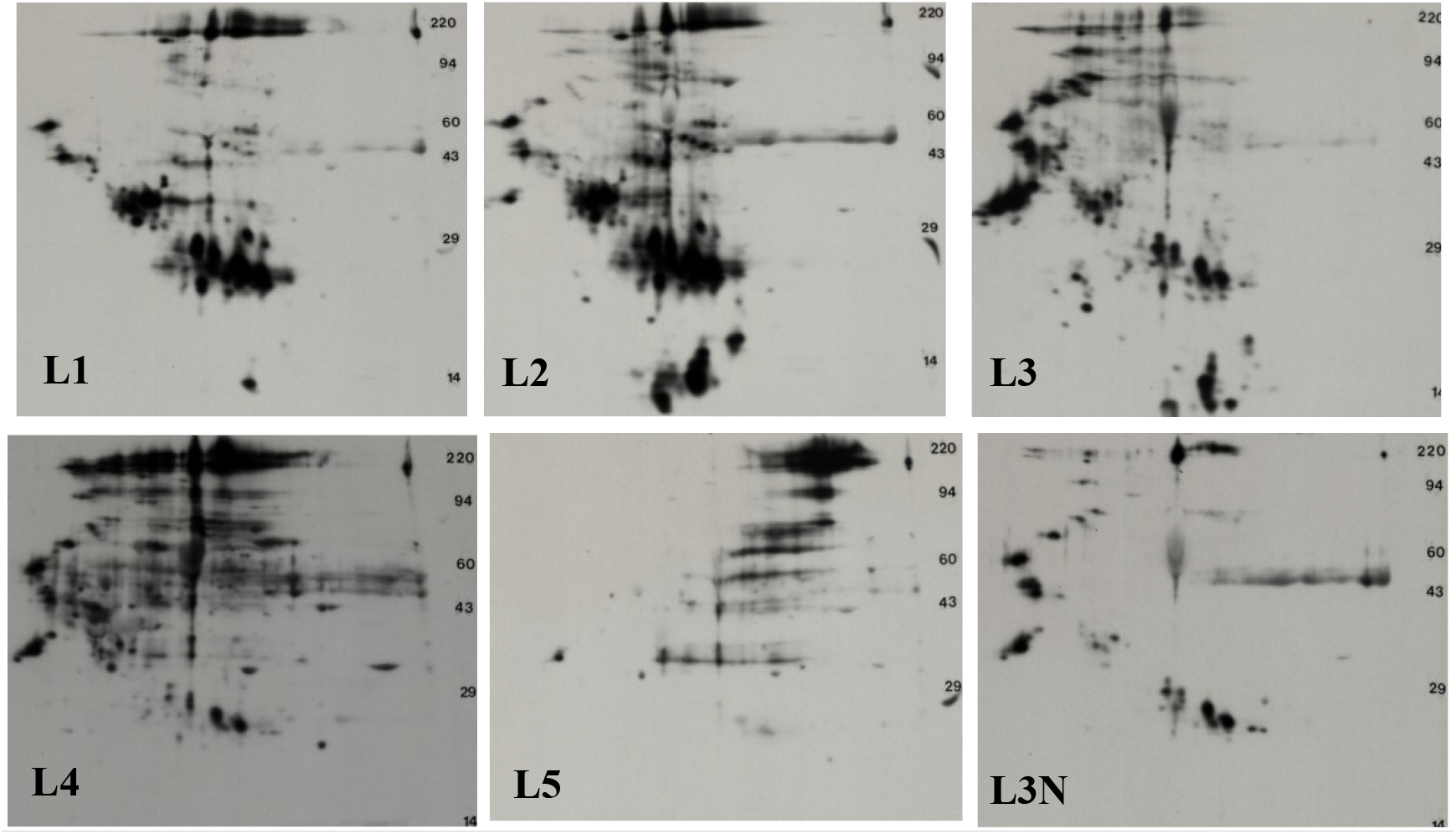
PDGFRB 2D WB patterns for LSCC samples L1-L5 along with L3N. The large amount of proteolysis visible on these images is suggestive of lysosomal digestion for PDGFRB in samples L1-L4 and L3N. In contrast, the PDGFRB in L5 shows lower MW breakdown products but does not show the diagonal pattern seen in the other five WBs. For L5, the high MW species, that co-migrates with a portion the pTyr signal, is the predominant form.

#### Tumor pTyr signal stability

Individual aliquots of L3 and L5 tumor homogenates prepared in SDS buffer and stored at -80° C were used as standards for Group 2 WB tests. They showed stable pTyr-RTK WB signals for approximately three years. Because of aliquot stability we anticipated that signal in solid tumor portions initially refrozen at -80° C and never rethawed would be stable as well. This was not the case. The remaining frozen half of L3 showed only trace pTyr-protein signal after three years (Supplemental Figure S3). An aliquot that had been thawed and refrozen also lost its signal.

## Discussion

NGS is used in precision oncology to identify genomic alterations (biomarkers) in tumor biopsies that predict cancer drug success. However, for LSCC, the predictions have been off the mark. For example, the FGFR1 gene, amplified in approximately 20% of LSCC patients, is an obvious biomarker candidate [30]. A small clinical trial of an FGFR1 inhibitor, dovitinib, targeted 26 prescreened LSCC patients with a median FGFR1 copy number of 6.8 (range 5-48 copies). However, after 16 months of treatment, only three patients had a partial response with durations of 5-6 months and none had a complete response. The FGFR copy numbers of the three responders were at the low end (5.5 – 5.8). FGFR mRNA was measured in 16 of the 26 patients. Of these, four showed no FGFR mRNA transcripts, and expression levels for the rest were not associated with either gene amplification or patient response [31, 32].

The large Lung-Map trial, which included six biomarker drugs targeted to 1,864 patients with advanced LSCC, also gave disappointing results. The median OS for the combined biomarker treatment group was shorter (5.9 mo) than that of the chemotherapy control group (7.7 mo) [33, 34].

A cancer driver gene is defined as one whose mutations increase net cell growth under the specific microenvironmental conditions that exist in the cell in vivo [35]. While RTKs are well-known cancer drivers they are not always directly activated by mutations. Some may become active when tumor suppressor genes (TSG) cease functioning because of genomic alterations. TSG mutations are recessive, so a single allele mutation does not cause carcinogenesis. But other inactivating mechanisms, epigenetic silencing for example, may contribute to loss of activity. Since there are many possible TSGs that might affect RTK activity, it is difficult to identify the culprits [36]. Prime candidates are the protein tyrosine phosphatases that regulate RTK tyrosine phosphorylation [37].

Another precision oncology problem is related to the central dogma: DNA makes RNA makes protein. Currently, gene mutation biomarkers are used to predict drug responses without consideration of transcription and translation. In fact, there is considerable evidence that mRNA levels correlate poorly with protein levels [38]. Wang et al. found 13,413 transcript and protein pairs in 29 healthy human tissues, with a median correlation between the transcriptome and proteome of 0.35. In 12% of the pairs, a negative correlation was observed, and some proteins could not be detected despite being highly expressed mRNAs [39]. One reason for this discrepancy is that at least 27% of human genes code for proteins that post-translationally modify or otherwise affect other proteins. (Supplemental Table 1.) As discussed by Roehri et al., proteome-based pathology is likely to be the next frontier in precision medicine [40].

Intratumor heterogeneity (ITH), caused by tumor subclones growing at different speeds is also a major issue. Jamal-Hanjani et al. tracked genomic changes in 327 tumor regions from 100 patients with lung cancer, including 32 patients with LSCC. More than 75% of the tumors carried subclonal driver alterations that varied from region to region within a tumor [41]. Ramifications have been discussed by Ramon et al. [42]. Although targeted monotherapy may cause transient regression by inhibiting a subset of tumor cells, the subsequent outgrowth of resistant subclones leads to recurrence.

Finally, RTKs have multiple ligands: EGFR has seven ligands capable of triggering dimerization and phosphorylation [17] and FGFR has 18 [43]. Because EGFR and FGFR proteins are often overexpressed in LSCC [30, 44], mutation-driven increases in any serum ligand would likely trigger subclone growth. In fact, an anti-EGF vaccine clinical trial for NSCLC showed that serum EGF is useful as a vaccine biomarker [45].

To circumvent these problems, we propose testing large (∼ 1 g) tumor biopsies using pTyr-RTK WB on a research basis to identify tumors with active pTyr-RTK drivers. Subsequent RNA-seq analysis of those tissues should reveal mRNA fingerprints that correlate with the known drivers and/or common pathways. Such mRNA fingerprints should predict drug success more accurately than any single entity. For example, Chang et al. studied the mRNA expression levels of housekeeping genes (HKG) in human lung cancers. They found that the mRNA fingerprints of 106 HKG allowed the separation of LADC (n=40) from LSCC tissues (n=18). They concluded that the mRNA expression patterns of HKG reflect the specific cellular state of the sample at the time of collection [46]. Jaakkola and Elo discussed nine other deconvolution methods for estimating cell type-specific differential expression in tissue samples [47].

Here, we describe the results of a feasibility test to determine if the first step, ECL WB, is useful for screening cancer tissue for the presence of pTyr-RTKs, that is, specific, actionable, driver proteins, including both mutated and WT species.

### Major Findings

We used a sensitive method of pTyr WB [48] to analyze homogenates of resected tumor tissue from 12 LSCC, 2 LADC, and 11 lung controls for the presence of RTKs activated by tyrosine transphosphorylation. Since RTKs are known to be high MW proteins, we used this parameter as the initial criterion for a positive result. Two of the 12 LSCC tumors (L3 and L5) showed strong pTyr WB bands near 220 kDa (Figures 1-5). Two other tumors, L4 and L11, showed pTyr bands darker than the controls at the same MW but lighter than L3 and L5. One LADC tumor with an activating EGFR exon 19 mutation (L8, Figure 3) showed weak EGFR and pTyr-EGFR signals compared to LSCC tumors L3, L4, and L5 run on the same WB, rather than the expected strong EGFR signal. This suggests that activating gene mutations detected by NGS do not necessarily predict increased protein activity. In total, our results (Figures 2 and 3) showed higher EGFR protein levels in tumors than in controls in 7/12 LSCC samples (L1, L3, L4, L5, L11, L12, and L13) or 58%, in agreement with IHC reports for LSCC [49].

Unequivocal identification of pTyr-RTKs by WB was hindered by glycosylation. RTKs are transmembrane proteins with branching N-glycan chains in their extracellular domains [50]. Glycans dramatically alter the protein MW [51] and end in negatively charged sialic acids that impart charge heterogeneity. Thus, the 2D spot outlines of the RTKs were diffuse. The heavy tumor protein glycosylation that we encountered is common in cancer [52] and could be a target for immunotherapy [53].

Initial attempts to directly identify low-abundance pTyr-RTK proteins by LC/MS/MS of protein spots cut from 2D gels (500 μg gel load) were unsuccessful (Supplemental Figure S1). Pattern matching of 2D pTyr WBs to different RTK protein WBs (Figures 5 and 6) suggested that a single driver, pTyr-EGFR, was active in L3, whereas two drivers, pTyr-EGFR and pTyr-PDGFRB, were simultaneously active in L5. For the latter, a strip-and-reprobe WB experiment, in where pTyr, EGFR, and PDGFRB antibodies were used on the same 2D gel, confirmed that both pTyr-RTKs were present in L5 (Figure 6).

Although PDGFRB is normally associated with mesenchymal tissue [54], Zhang et. al. observed PDGFB and PDGFRB in 74% and 78% of tumor cells, respectively, in 66 patients with LSCC using IHC [26]. Given the different breakdown patterns of PDGFRB in L5 vs. L1-L4 and L3-NAT (Figure 7) and the overlap of the anti-PDGFRB WB with a large portion of the pTyr WB pattern (Figure 6), we concluded that this RTK was active in the L5 tumor.

Minor pTyr 2D spots were detected in L3 and L5 tumors at a lower MW (red arrows, Figure 5), which did not match any breakdown products in the EGFR or PDGFRB patterns. These could be additional cancer drivers, such as pTyr-FGFR1 or pTyr-DDR2 in minor subclones.

### Additional Findings, Existing Literature

Other groups identified pTyr-EGFR in lung tumor biopsies using IHC with antibodies specific for one of the six pTyr-EGFR docking sites: pTyr-845, -1045, -1068, -1086, -1148, and -1173 [55]. Kanematsu et al. showed that phosphorylation, but not EGFR overexpression, is associated with poor prognosis of NSCLC using an antibody against pTyr845 [56]. Wang et al. used antibodies against pTyr1068 and pTyr1173 in biopsies from stage III/IV NSCLC patients treated with erlotinib or gefitinib. For the latter, the drug response of 84 patients with WT EGFR varied with the phosphorylation status of the two sites [20].

Rikova et. al. used IHC, pTyr WB, and MS phosphoproteomics to characterize 150 NSCLC tumor samples, including 60 LSCC samples. Results were mostly unsorted regarding histology, but one table shows MS data from 12 patients with LSCC. For the latter, pTyr-DDR1 and pTyr-EGFR predominated in five tumors, pTyr-EGFR predominated in two tumors, pTyr-ALK predominated in one, pTyr-Met predominated in one, and pTyr-PDGFRA predominated in three. Ten pTyr-RTKs were identified in 12 LSCC patients [29].

Marked differences in sample preparation exist between MS and WB, which could affect reported findings. For example, the detergent SDS, which dissolves membrane proteins but interferes with MS ionization, is often omitted from the MS sample buffers. Rikova et al. homogenized tumor tissues in urea buffer, followed by centrifugation to remove particulates. The discarded pellets probably contained membrane bound RTKs. Because MS requires trypsin digestion prior to analysis, it cannot distinguish between peptides from intact RTKs and those from RTKs degraded after endocytosis. The latter is a major regulator of RTK signaling [29].

### Study Limitations

The pTyr-RTK proteins found in L3 and L5 tumors by 1D WB screening proved surprisingly difficult to identify by 2D WB. MS identification of proteins cut from stained 2D gels is usually straightforward [57], but in this case, not enough material was present for MS. (Supplemental Figure 1). The concentration of pTyr-peptides using immunoaffinity beads, as described by Rikova et al. [27] was not feasible. While MS is a key method for unknown protein identification, it is difficult to standardize [58]. Identification of tyrosine phosphorylation sites can be problematic [59].

The pictorial presentation of 2D results became a problem. Comparing pTyr WB results for a few samples at the onset was straightforward; however, it was difficult to keep track of images from multiple tumors and antibodies. A way to express results numerically is needed to compare the results over time and between laboratories. To that end, our lab changed focus to generate an internal pTyr-protein standard that could be added to every sample for normalization. A kinase transphosphorylation reaction was performed in vitro using a 48 kDa recombinant fragment containing the kinase domain of a commercial anaplastic lymphoma kinase (ALK). The usefulness of the standard has been verified and published [48]. For future work, adding a known amount of pTyr-ALK48 (or similar phosphoprotein) to every sample would allow the numerical expression of results as a spot density ratio.

### Future Directions

In future studies, unequivocal identification of pTyr-RTK signals could be achieved using antibodies against specific pTyr-EGFR residues, such as those used by Wang et al. [20] for IHC. Specific pTyr-residue antibodies for rare RTK candidates should be generated and validated. Immunoprecipitation of intact pTyr-proteins from large biopsy samples might provide enough material for confirmation of pTyr-RTK identity by orthogonal methods, pTyr-WB and MS.

Immunotherapy and/or chemotherapy are usually the first-line treatments for LSCC, as shown in Keynote 407 [60] and ChecMate227 [61]. However, what will the second-line treatment be when immunotherapy fails? As discussed by Santos and Rodriguez, the RTK inhibitors ramucirumab (VEGFR-2 inhibitor) and afatinib are candidates for second-line treatment [62]. Developing methods to predict which patients will respond to these inhibitors is an important goal.

### Overall Conclusion, Major Impact

Precision medicine strategies for drug delivery are often complex [63]. Knowledge of whether actionable pTyr-RTKs, either WT or mutant, are present in a tumor mass would help inform treatment decisions. Results of this feasibility experiment show that pTyr WB is sensitive enough to screen LSCC tissue biopsies to find subsets with active pTyr-RTK(s) protein drivers, especially pTyr-EGFR and pTyr-PDGFRB, but also other RTKs.

Flash-frozen tumor tissue is seldom available from patients; therefore, the WB approach will likely not be a diagnostic test. However, WB could be used to identify LSCC research samples containing single and multiple pTyr-RTK drivers, and perhaps other oncoproteins. Such knowledge could be used to inform genomic testing, especially RNA-seq, to improve biomarker predictions. Our results need to be confirmed in a larger number of human LSCC samples using an internal pTyr standard for semi-quantitative results, followed by RNA-seq analysis. Using samples from rapid autopsies [64] would allow primary tumors to be compared with metastatic sites to assess differences.

Rodriguez and Pennington proposed that collaborative proteogenomics, including multi-omic approaches, might be a future direction for precision oncology [65]. We are convinced that this is the case.

## Materials & Methods

### Sample preparation

Fourteen human lung tumor and 11 control lung samples were purchased in two groups from a human biobank (ILS Bio, LLC, now BioIVT) Chesterfield, MD. Under the definition of human research subject [45 CFR4 46.102(f)], the Office of Human Resource Protection does not consider research involving only coded private information or biospecimens (retrospective or remnant) to involve human subjects. These specimens were not specifically collected for the proposed research project. Because investigators cannot readily ascertain the identity of the individuals to whom the coded specimens pertain, the requirement for documented informed consent is not applicable.

The details of the samples are listed in Table 1. All samples were prepared on ice using ice-cold reagents. Tumor samples were cut in half with a scalpel; half of each tumor was frozen on dry ice and stored at -80°C in a monitored freezer without intermediate thawing. The other half was rinsed with ice-cold Tris-buffered saline TBS) [20 mM Tris (MP Biomedicals, Solon, OH), 500 mM NaCl (ThermoFisher, Waltham, MA), pH 7.5], and then placed in a motorized glass-Teflon homogenizer. The tumor tissue was homogenized on ice with Osmotic Lysis Buffer [10mM Tris, 0.3% SDS (IBI Scientific, IA), pH 7.4)], Protease Inhibitor stock [0.2 mM AEBSF (Calbiochem, Burlington, MA), 10 μg/ml leupeptin (Sigma-Aldrich, St. Louis, MO), 3.6 μg/ml E-64 (Sigma-Aldrich), 5 mM EDTA (Calbiochem), and 56 μg/ml benzamidine (Sigma-Aldrich)], Phosphtase Inhibitors I and II (MilliporeSigma, Burlington, MA), Nuclease stock [5 U/ml RNase (Sigma-Aldrich) and 150 U/ml DNase (Sigma-Aldrich), 5 mM MgCl_2_ (Sigma-Aldrich), 10 mM Tris, pH 7.0], and SDS buffer (5% SDS, 10% glycerol (Sigma-Aldrich), 62.5 mM Tris, pH 6.8).

The tube containing the homogenized sample was placed in a boiling water bath for 5 min, and the protein concentration was determined using the BCA assay (ThermoFisher). Finally, each sample was diluted to 4 mg/ml in SDS buffer containing 5% β-mercaptoethanol. Aliquots were stored at -80°C.

## 1D SDS PAGE

SDS slab gel electrophoresis was carried out in hand-poured 10% acrylamide slab gels (13 × 15 cm, 0.75 mm thick), as described previously for the second dimension of 2DE [48].

Electrophoresis was performed for approximately 4 h at 15 mA/gel. The following proteins (MilliporeSigma) were used as MW standards in one lane on every gel: myosin (220,000), phosphorylase A (94,000), catalase (60,000), actin (43,000), carbonic anhydrase (29,000), and lysozyme (14,000).

## 2D SDS PAGE

Two-dimensional SDS-PAGE was performed as previously described [48]. Briefly, isoelectric focusing was performed in polymerized polyacrylamide tube gels (130 mm long x 2.3 mm internal diameter) that were sealed at the bottom with parafilm and poured using 2% ampholines (Serva Electrophoresis, Heidelberg, Germany). Ampholines were pH 3-10 Iso-Dalt or a 1:1 mixture of pH 4-6 Ampholines and Servalyte pH 5-8. Samples were loaded at the top (basic end) of the polymerized acrylamide tube gel, and isoelectric focusing was performed at 9600 volt-hrs.

Tube gels were extruded under water pressure and equilibrated for 10 min in buffer “O” (10% glycerol, 50 mM dithiothreitol, 2.3% SDS, and 0.0625 M Tris, pH 6.8). The equilibrated tube gels were frozen on dry ice to prevent protein diffusion and thawed immediately before loading onto the 2^nd^ dimension slab gel. Each tube gel was sealed in 1 ml of agarose to the top of a stacking gel overlaying a 10% acrylamide slab gel, and electrophoresis was performed for approximately 4 h at 15 mA/gel. The same MW standards used for 1D SDS PAGE were loaded into an agarose well at the basic end of the tube gel. One microgram of tropomyosin was loaded with each sample on the IEF tube gel as an internal standard.

### Western Blotting

After slab gel electrophoresis, the gels were placed in transfer buffer (10 mM CAPS, pH 11.0, 10% methanol) and transferred onto PVDF membranes overnight at 200 mA and approximately 100 volts/two gels. The blots were stained with Coomassie Brilliant Blue R-250 (Sigma-Aldrich) and scanned.

Coomassie blue-stained PVDF membranes were wet in 100% methanol to remove the stain, rinsed in tween-20 (Bio-Rad) TBS (TTBS), and blocked for two hours in 5% bovine serum albumin diluted in TTBS. Primary antibody incubations were performed overnight on an orbital shaker in 2% bovine serum albumin using the antibody dilutions shown in Table 2. Blots were rinsed 3 × 10 min in TTBS and incubated with a secondary antibody diluted 1:2000 for 2 h. After rinsing again as above, the blots were treated with Pierce ECL reagent (ThermoFisher) and exposed to X-ray film [Kodak BioMax MR film (ThermoFisher) or GE Amersham Hyperfilm ECL)], followed by film development with an automatic Konica Minolta Medical Film Processor SRX-101A. Stripping of 2D WBs was performed as described by GE Healthcare ECL Advance WB Detection Kit Product Booklet, Code RPN2135, before re-probing with a second primary antibody. GE Amersham Hyperfilm was used for overlay images. The antibodies that were used are listed in Table 2.

**Table 2.** Antibodies used for WBs. *For GE Hyperfilm, primary antibodies were diluted 1:5000 or 1;10,000.

| Antibody | Supplier | Source | Dilution | Secondary Antibody |
| --- | --- | --- | --- | --- |
| Phosphotyrosine [66] | ExAlpha, Clone PY20<br>Cat # X1021 | Mouse | 1:1000 or<br>1:5000* | Anti-mouse IgG HRP<br>from sheep |
| EGFR | Cell Signaling<br>Cat # 4267S | Rabbit | 1:1000 or<br>1:10,000* | Anti-rabbit IgG HRP<br>from sheep |
| PDGFRB | Cell Signaling<br>Cat # 4564S | Rabbit | 1:10,000* | Anti-rabbit IgG HRP<br>from sheep |
| VEGFR 2 | Cell Signaling<br>Cat # 2479S | Rabbit | 1:10,000* | Anti-rabbit IgG HRP<br>from sheep |
| MET | Cell Signaling<br>Cat # 3148S | Mouse | 1:1000 | Anti-mouse IgG HRP<br>from sheep |
| ALK | Cell Signaling<br>Cat # 3791S | Mouse | 1:1000 | Anti-mouse IgG HRP<br>from sheep |

## Acknowledgments

The authors thank Judith Finlay and Lawrence Grossman for stimulating discussions and critical comments on the manuscript.

