## Supplemental Figures 1-3, Supplemental Table 1 for "Detection of Activated Receptor Tyrosine Kinases in Human Lung Squamous Cell Carcinoma"

### Figure S1

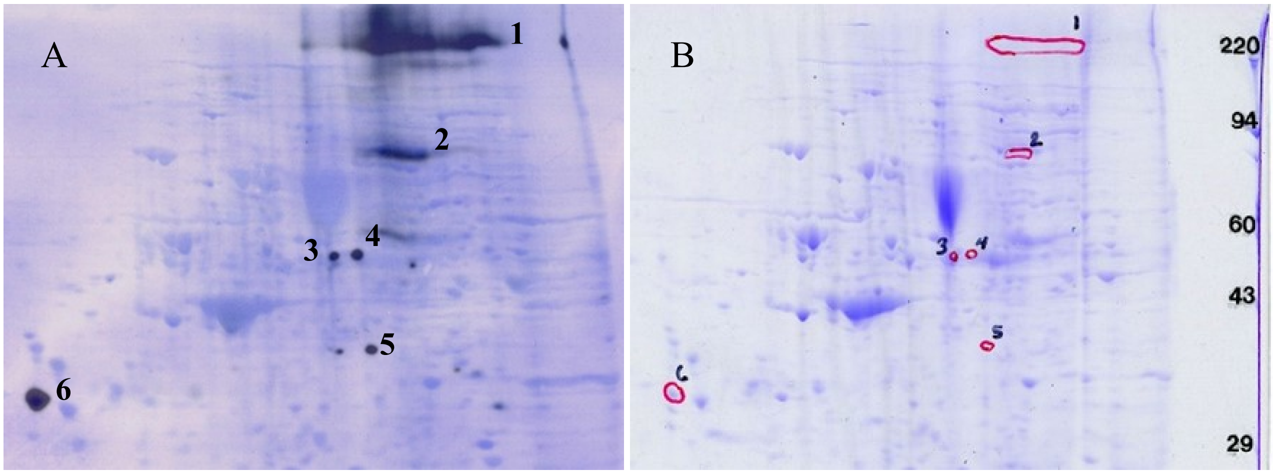


| Kendrick #1 (KL_XS_002822_002) | | | | |
|---|---|---|---|---|
| Protein Identified | Mascot Score | # Peptide sequences Identified | NCBI or SwissProt Accession # | MW |
| 90kDa heat shock protein [Homo sapiens] | 126 | 12 | gi\|306891 | 83,668 |
| Human Transglutaminase 2 Complex With Adenosine 5' Triphospha | 54 | 4 | gi\|303324982 | 71,551 |
| filamin A [Homo sapiens] | 42 | 9 | gi\|53791219 | 280,742 |

| Kendrick #2 (KL_XS_002822_005) | | | | |
|---|---|---|---|---|
| Protein Identified | Mascot Score | # Peptide sequences Identified | NCBI or SwissProt Accession # | MW |
| immunoglobulin mu heavy chain, partial [Homo sapiens] | 57 | 6 | gi\|390994295 | 67,003 |
| fatty acid synthase; FAS; multifunctional protein [Homo sapiens] | 45 | 9 | gi\|1049053 | 276,198 |

| Kendrick #3 (KL_XS_002822_008) | | | | |
|---|---|---|---|---|
| Protein Identified | Mascot Score | # Peptide sequences Identified | NCBI or SwissProt Accession # | MW |
| beta-tubulin [Homo sapiens] | 274 | 34 | gi\|1297274 | 51,054 |
| Aldehyde dehydrogenase, mitochondrial isoform 1 precursor [Homo sapiens] | 160 | 26 | gi\|25777732 | 56,985 |
| selenium-binding protein 1 isoform 1 [Homo sapiens] | 141 | 10 | gi\|16306550 | 53,068 |

| Kendrick #4 (KL_XS_002822_011) | | | | |
|---|---|---|---|---|
| Protein Identified | Mascot Score | # Peptide sequences Identified | NCBI or SwissProt Accession # | MW |
| beta-tubulin [Homo sapiens] | 327 | 36 | gi\|1297274 | 51,054 |
| selenium-binding protein 1 isoform 1 [Homo sapiens] | 84 | 7 | gi\|16306550 | 53,068 |

| Kendrick #5 (KL_XS_002822_014) | | | | |
|---|---|---|---|---|
| Protein Identified | Mascot Score | # Peptide sequences Identified | NCBI or SwissProt Accession # | MW |
| Lamin A/C transcript variant 1 [Homo sapiens] | 98 | 14 | gi\|57014047 | 74,381 |

| Kendrick #6 (KL_XS_002822_017) | | | | |
|---|---|---|---|---|
| Protein Identified | Mascot Score | # Peptide sequences Identified | NCBI or SwissProt Accession # | MW |
| tumor protein p53 binding protein 1 [Homo sapiens] | 41 | 10 | gi\|193784748/KAI4057465.1 | 43,578 |

Figure S1. NanoC-MS/MS analysis of six pTyr 2D protein spots in tumor L5. A. Overlay image showing six 2D pTyr WB protein spots aligned with the PVDF Coomassie blue image. B. Coomassie blue stained 2D gel from same run showing red outlines of spots cut for the MS analysis. The table below shows MS results. No RTKs were detected by nanoC-MS/MS within the spot 1 outline, likely because the high MW RTK protein was diffuse and in low abundance.

### MS Methods

***Protein digestion and peptide extraction.*** Proteins that were separated by SDS-PAGE/2D-PAGE and stained by Coomassie dye were excised, washed and the proteins from the gel were treated according to published protocols[1-5]. Briefly, the gel pieces were washed in high purity, high performance liquid chromatography (HPLC) grade water, dehydrated, cut into small pieces, destained by incubating in 50 mM ammonium bicarbonate, 50 mM ammonium bicarbonate/50% acetonitrile (A and then 100% acetonitrile under moderate shaking, followed by drying in a speed-vac concentrator. The gel bands were then rehydrated with 50 mM ammonium bicarbonate. The procedure was repeated twice. The gel bands were then rehydrated in 50 mM ammonium bicarbonate containing 10 mM DTT and incubated at 56ºC for 45 minutes. The DTT solution was then replaced by 50 mM ammonium bicarbonate containing 100 mM Iodoacetamide for 45 minutes in the dark, with occasional vortexing. The gel pieces were then re-incubated in 50 mM ammonium bicarbonate/50% acetonitrile, and 100% acetonitrile under moderate shaking, followed by drying in speed-vac concentrator. The dry gel pieces were then rehydrated using 50 mM ammonium bicarbonate containing 10 ng/µL trypsin and incubated overnight at 37ºC under low shaking. The resulting peptides were extracted twice with 5% formic acid/50 mM ammonium bicarbonate/50% acetonitrile and once with 100% acetonitrile under moderate shaking. Peptide mixture was then dried in a speed-vac, solubilized in 20 µL of 0.1% formic acid /2% acetonitrile.

***NanoLC-MS/MS.*** The peptide mixture was analyzed by reverse-phase nanoliquid chromatography (LC) and MS (LC-MS/MS) using a NanoAcuity UPLC (Micromass/Waters, Milford, MA) coupled to a Q-TOF Xevo G2 mass spectrometer (Micromass/Waters, Milford, MA), according to published procedures[1-5]. Briefly, the peptides were loaded onto a 100 µm x 10 mm NanoAquity BEH130 C18 1.7 µm UPLC column (Waters, Milford, MA) and eluted over a 60 minute gradient of 2-80% organic solvent (ACN containing 0.1% FA) at a flow rate of 400 nL/min. The aqueous solvent was 0.1% FA in HPLC water. The column was coupled to a Picotip Emitter Silicatipnano-electrospray needle (New Objective, Woburn, MA). MS data acquisition involved survey MS scans and automatic data dependent analysis (DDA) of the top six ions with the highest intensity ions with the charge of 2+, 3+ or 4+ ions. The MS/MS was triggered when the MS signal intensity exceeded 250 counts/second. In survey MS scans, the three most intense peaks were selected for collision-induced dissociation (CID) and fragmented until the total MS/MS ion counts reached 10,000 or for up to 6 seconds each. The entire procedure used was previously described[1-5]. Calibration was performed for both precursor and product ions using 1 pmolGluFib(Glu1-Fibrinopeptide B) standard peptide with the sequence EGVNDNEEGFFSAR and the monoisotopic doubly-charged peak with m/z of 785.84.

***Data processing and protein identification.*** The raw data were processed using ProteinLynx Global Server (PLGS, version 2.4) software as previously described[1-5]. The following parameters were used: background subtraction of polynomial order 5 adaptive with a threshold of 30%, two smoothings with a window of three channels in Savitzky-Golay mode and centroid calculation of top 80% of peaks based on a minimum peak width of 4 channels at half height. The resulting pk1 files were submitted for database search and protein identification to the in-house Mascot server (www.matrixscience.com, Matrix Science, London, UK) for database search using the following parameters: databases from **NCBI Human,** parent mass error of 0.5 Da with 1 $^{13}$C, product ion error of 0.8 Da, enzyme used: trypsin, three missed cleavages, propionamide as cysteine fixed modification and methionine oxidized as variable modification. To identify the false negative results, we used additional parameters such as different databases or organisms, a narrower error window for the parent mass error (1.2 and then 0.2 Da) and for the product ion error (0.6 Da), and up to two missed cleavage sites for trypsin. In addition, the pkl files were also searched against in-house ProteinLynx Global Server database version 2.4 (www.waters.com) using searching parameters similar to the ones used for Mascot search. The Mascot and PLGS database search provided a list of proteins for each gel band. To eliminate false positive results, for the proteins identified by either one peptide or a mascot score lower than 25, we verified the MS/MS spectra that led to identification of a protein.

### Figure S2

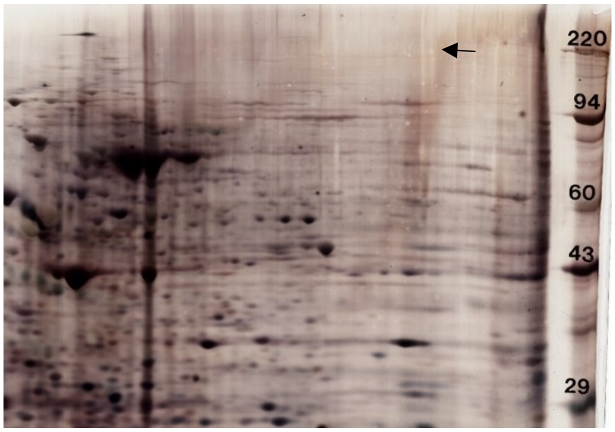


Figure S2. Silver stained 2D gel loaded with 50 ug tumor L5. The area where the high molecular weight pTyr WB 2D spot runs (arrow) shows only a faint haze.

### Figure S3

Figure S3. Variation of pTyr 2D WB signal with storage conditions for tumor L3. **A**. WB signal from an L3 aliquot stored for 17 months at -80 C (2763#3). **B**. Corresponding WB signal from the L3 tissue portion that had been re-frozen at -80 for three years before preparation in SDS buffer (2913#2). Virtually all pTyr WB signal was lost. **C.** WB signal from an L3 aliquot that had thawed, an aliquot removed, then refrozen at -80 C for one month rather than being discarded. (2912#4). Film exposure times: 15 min for A; 10 min for B and C.

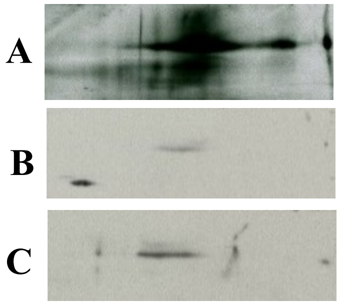

### Table S1

| Protein families that perform post-translational modifications (PTM) or actions (TFs &RBPs). | Number |
|---|---|
| Transcription factors (TF) [6] | 1,639 |
| RNA-binding proteins (RBP) [7] | 1,542 |
| Tyrosine kinases (58 RTK & 32 TK) [8] | 90 |
| Tyrosine phosphatases [9] | 107 |
| Serine/threonine kinases [8] | 418 |
| Serine/threonine phosphatases [10] | 30 |
| Proteases (PTM) [11] | 588 |
| Glycosyltransferases (PTM) [12] | 244 |
| E3 ubiquitin ligases (PTM) [13] | 377 |
| Glutathionine S-transferases [14] | 16 |
| Sulfotransferases [15] | 13 |
| Histone lysine methyltransferases [16] | 50 |
| Histone acetyltransferases and deacetylases [17] | 35 |
| DHHC3 Palmitoyltransferases [18] | 23 |
| 2-OG-dependent dioxygenases [19] | 70 |
| Total | 5,192 |
| Total protein-coding genes in human genome [20] | 19,116 |

Table S1. Protein families that act post-translationally via expression/repression of protein transcripts (TFs), translation of mRNA transcripts into proteins (RBPs), or via post-translational modifications (PTMs). Genes in these families make up at least 27% of total protein-coding genes in the human genome. Gene mutations of some of these proteins likely exacerbate cancer indirectly, but which ones and how many cannot yet be predicted.
